# Sortilin-related receptor is a druggable therapeutic target in breast cancer

**DOI:** 10.1101/2021.03.09.434556

**Authors:** Hussein Al-Akhrass, Mika Pietilä, Johanna Lilja, Ella-Maria Vesilahti, Johanna M. Anttila, Heidi M. Haikala, Pauliina M. Munne, Juha Klefström, Emilia Peuhu, Johanna Ivaska

## Abstract

In breast cancer, the currently approved anti-receptor tyrosine-protein kinase erbB-2 (HER2) therapies do not fully meet the expected clinical goals due to therapy resistance. Identifying alternative HER2-related therapeutic targets could offer means to overcome these resistance mechanisms. We have previously demonstrated that an endosomal sorting protein, sortilin-related receptor (SorLA), regulates the traffic and signaling of HER2 and HER3, thus promoting resistance to HER2-targeted therapy in breast cancer. This study aims to assess the feasibility of targeting SorLA using a monoclonal antibody. Our results demonstrate that anti-SorLA antibody (SorLA ab) alters the resistance of breast cancer cells to HER2 monoclonal antibody trastuzumab *in vitro* and *in ovo*. We found that SorLA ab and trastuzumab combination therapy also inhibits tumor cell proliferation and tumor cellularity in a mouse xenograft model of HER2-positive breast cancer. In addition, SorLA ab inhibits the proliferation of breast cancer patient-derived explant three-dimensional cultures. These results provide for the first time proof-of-principle that SorLA is a druggable target in breast cancer.

## Introduction

The receptor tyrosine-protein kinase erbB-2 (HER2) belongs to the HER family of cell surface receptors, which transduce extracellular cues into intracellular signals upon receptor homo- or heterodimerization (1). The amplification of the gene encoding HER2 occurs in 15-30% of breast tumors defining a histopathological breast cancer subtype. The diagnosis of HER2-positive breast tumors guides therapy decisions with anti-HER2 therapeutics dramatically improving patients’ clinical outcome (1,2). However, HER2-targeted therapies fail in achieving durable efficacy due to acquired resistance leading to distant-organ metastases in the most challenging clinical setting (3,4). Currently, there are no clear treatment options for patients who have progressed after two lines of anti-HER2 therapy, with reported response rates being extremely poor (ranging from 10 % to 22 %) (5,6). This indicates a highly unmet medical need for patients progressing under current anti-HER therapies. In the published literature, several therapy-resistance mechanisms have been reported, most notably sustained oncogenic signaling that compensates for HER2 inhibition through altered expression of receptor tyrosine kinases such as HER3 (7,8). HER2 interacts with HER3 forming the most signaling-potent dimer among the HER family (9). Therefore, targeting HER3 may alleviate anti-HER2-therapy resistance in breast cancer; however, this approach appears to be extremely challenging despite the extensive preclinical and clinical efforts due to HER3 containing a pseudokinase domain in its intracellular region which renders it undruggable with kinase inhibitors (10,11).

Sortilin-related receptor (SorLA) is an intracellular sorting protein and a member of the family of vacuolar protein sorting 10 protein (VPS10P)-domain receptors (12). The N-terminal SorLA extracellular/luminal part contains multiple subdomains shown to mediate ligand binding/discharge, while the short C-terminal tail, containing trafficking signals, binds to cytosolic adaptor proteins to assemble protein complexes orchestrating SorLA traffic (13–15). The mature SorLA protein resides mainly in the trans-Golgi network and undergoes constitutive antero- and retrograde trafficking to the plasma membrane through endosomes. Due to its central role in protein trafficking, SorLA has been implicated in the development and/or progression of neurological and metabolic diseases, and most recently in cancer (16–19).

Our previous studies unraveled a HER2-therapy resistance mechanism in which SorLA supports HER2 protein levels and oncogenicity in breast and bladder cancer *in vitro* and *in vivo* (16). In addition, we have demonstrated that SorLA promotes HER2-HER3 endosomal recycling to sustain their oncogenic signaling *in vitro* as well as in an anti-HER2 therapy insensitive brain xenograft *in vivo* model (19). The present study outlines the translational extension of our previous findings, and aims to assess the druggability of SorLA in breast cancer. Our results demonstrate that an anti-SorLA antibody (SorLA ab) alters the resistance of metastatic breast cancer cells to the HER2 monoclonal antibody trastuzumab *in vitro* and in chick chorioallantoic membrane (CAM) assays. In a HER2-positive breast cancer mouse xenograft model, SorLA ab and trastuzumab combination treatment inhibits tumor cell proliferation and tumor cellularity. In addition, SorLA ab may be clinically relevant since it inhibits the proliferation of patient-derived HER2-amplified breast explant cultures. These results demonstrate for the first time that SorLA is an achievable therapeutic target in breast cancer. The implications of this proof-of-principle SorLA druggability study may be relevant for SorLA-promoted human cancer, beyond HER2-positive breast cancer.

## Materials and Methods

### Cell culture and reagents

BT-474 cells (ATCC, HTB-20) were grown in RPMI-1640 (Sigma-Aldrich, R5886) supplemented with 10% fetal bovine serum (FBS; Sigma-Aldrich, F7524), 1% vol/vol penicillin/streptomycin (Sigma-Aldrich, P0781-100ML) and L-glutamine (Sigma-Aldrich, G7513-100ML). MDA-MB-361 cells (ATCC, HTB-27) were grown in Dulbecco’s modified essential medium (DMEM; Sigma-Aldrich, D5769) supplemented with 20% FBS, 1% vol/vol penicillin/streptomycin and L-glutamine. SK-BR-3 cells (ATCC, HTB-30) were grown in McCoy’s 5A Medium, Modified with L-glutamine (Sigma-Aldrich, M9309) supplemented with 10% fetal bovine serum and 1% vol/vol penicillin/streptomycin. All cells were cultured in a humidified incubator set at 5% CO2 and 37 °C. All cells were tested bimonthly for mycoplasma using MycoAlert™ mycoplasma detection kit (Lonza, #LT07-418) and MycoAlert™ assay control set (#LT07-518). Cell lines were not independently authenticated for this study. The antibodies used are described in Supplementary Table 1.

### Western blot

Cells were washed with ice-cold Dulbecco’s phosphate-buffered saline (DPBS, Gibco™, 11590476) prior to lysis with cell lysis buffer (CST, #9803) supplemented with 1% protease/phosphatase inhibitor cocktail (CST, #5872). Lysates from the MMTV/C-neu transgenic mice (20) were generously provided by Jukka Westermarck (Turku Bioscience Centre, (21)). Cell lysates were sonicated and cleared by centrifugation at 18,000 g for 10 min. 30 μg of cleared lysates were subjected to SDS-PAGE under denaturing conditions (4–20% Mini-PROTEAN TGX Gels) and were transferred to nitrocellulose membranes (Bio-Rad Laboratories). Membranes were blocked with Odyssey Blocking Buffer (LI-COR Biosciences, #927-40000) and incubated with the indicated primary antibodies, diluted in blocking buffer and PBS, overnight at +4 °C. Membranes were then washed three times with TBST (Trisbuffered saline and 0.1% Tween 20) and incubated with fluorophore-conjugated secondary antibodies diluted (1:2000) in blocking buffer at room temperature for 1 h. Membranes were scanned using an infrared imaging system (Odyssey; LI-COR Biosciences). The following secondary antibodies were used: donkey anti-mouse IRDye 800CW (LI-COR, 926-32212), donkey anti-mouse IRDye 680RD (LI-COR, 926-68072), donkey anti-rabbit IRDye 800CW (LI-COR, 926-32213) and donkey anti-rabbit IRDye 680RD (LI-COR, 926-68073). The band intensity of each target was quantified using ImageJ (NIH) (22) and normalized to the loading control band intensity in each lane.

### Transient siRNA-mediated knock down

Cells were transfected 72 h before experiments using Lipofectamine RNAiMAX reagent (Invitrogen, P/N 56532) according to the manufacturer’s instructions. SORL1-targeting siRNAs were obtained from Dharmacon – siSORL1 #3 (J-004722-07, (5’CCGAAGAGCUUGACUACUU3’)), siSORLA #4 (J-004722-05, (5’CCACGUGUCUGCCCAAUUA3’)). Allstars (Qiagen, 1027281) was used as a negative control. All siRNAs were used at a final concentration of 20 nM.

### Cell viability assays

Cells were silenced for SorLA in 6-well plates and then replated on 96-well plates (5,000 cells/well) in a volume of 100 μL and allowed to grow for 72 h. After experiments, 10 μL/well of WST-8 (cell counting kit 8, Sigma-Aldrich, 96992) reagent was added. After 3 h of incubation at 37 °C, 5% CO2, absorbance was read at 450 nm (Thermo, Multiscan Ascent). Medium without cells was used as a background control, subtracting this from the sample absorbance readings. Cell viability was calculated as a ratio of endpoint absorbance relative to control cells.

### Soft agar assay

Bottom agar (1.2 %, prepared in normal medium) was casted at the bottom of 12-well plates. BT-474 cells stably expressing green fluorescent protein (GFP) with either control shRNA or SORL1 shRNA were resuspended at a concentration of 20 000 cells/1.5 ml in agar (top agar; 0.4 % prepared in normal medium) and seeded on top of the pre-casted bottom agar. Plates were shortly incubated at +4°C to accelerate the solidification of top agar, thus maintaining the single cell suspension and preventing cells from dropping down to the border between the top and bottom agar layers. After top agar was solidified, 1 ml of normal medium was added on top. Medium was changed once a week. Soft agar colony size was measured after 5 weeks of growth by imaging the GFP signal on a fluorescence microscope and measuring the area of GFP-positive colonies using Image J (NIH) (22).

### Colony formation assay

MDA-MB-361 and BT-474 expressing either control shRNA or SORL1 shRNA were seeded on 6-well plates (1000 cells/well). Medium was changed twice a week and after 4 weeks of growth colony number was assessed by manual counting of colonies with more than 10 cells.

### Cell cycle analysis

BT-474 cells transfected with control or SorLA-targeting siRNA were trypsinized, harvested using centrifugation and suspended in ice cold PBS (1ml). Cells were fixed with 70 % EtOH (ice cold) added in a dropwise manner with gentle vortexing of cells. Cells were then stored at +4°C until propidium iodide (PI; Sigma-Aldrich, #P4864-10ML) staining was performed. Cells were washed twice with PBS, suspended in PI solution (25 μg/ml PI solution in PBS with RNAase A; MACHEREY-NAGEL, #740505). and incubated for 10 minutes on ice. Samples were protected from light and kept on ice until flow cytometry analysis. Samples were run on an LSR II flow cytometer and cell cycle analysis was performed with FlowJo (BD Biosciences).

### Chorioallantoic membrane (CAM) assay

Fertilized chicken eggs were washed with 70 % EtOH and the development was started by placing the eggs in a 37 °C incubator with 50-60% moisture (egg developmental day 0 (EDD0)). On EDD3, a small hole was made in the eggshell to drop the CAM. On EDD10, the egg shell was opened and a plastic ring was placed on the CAM and one million control- or SORL1 siRNA-transfected cells were implanted inside the ring in 20 μl of 50% Matrigel. On EDD16, tumors were imaged, dissected and weighed.

For CAM treatment experiments, the egg shell was opened and a plastic ring was placed on the CAM on EDD7. One million MDA-MB-361 cells, with either IgG-control, anti-SorLA antibody, trastuzumab or trastuzumab in combination with anti-SorLA antibody (customized without azide, and with low endotoxin levels) were implanted inside the ring in 20 μl of 50% Matrigel. On EDD10, the antibody treatments were repeated by pipetting antibody solutions on tumors inside the ring. Tumors were dissected and weighed on EDD12.

### Orthotopic *in vivo* xenografts

All animal studies were ethically performed and authorized by the National Animal Experiment Board and in accordance with The Finnish Act on Animal Experimentation (Animal license number ESAVI-9339-04.10.07-2016). For orthotopic inoculation, MDA-MB-361 cells were suspended in PBS with 50% Matrigel. Cells (5 × 10^6^ in 50 μl) were injected orthotopically into the abdominal mammary gland fat pad of 7-8 week-old ovariectomized Nude female mice (Hsd:Athymic Nude-foxn1nu, Envigo, France) under isoflurane anesthesia. For estrogen supplementation, a 60-day release E2 (1.5 μg/day) MedRod implant (PreclinApps Ltd) was inserted s.c. at the flank of the mice. Therapies were administrated (i.p.) twice a week from day 15 post-engraftment when the average size of tumors had reached 100 mm^3^. Mice were blindly randomized into three groups: a control group receiving a combination of lgG1 (1 mg/kg) and lgG2a (10 mg/kg), a monotreatment group receiving trastuzumab (1 mg/kg), or a combination-treatment group receiving trastuzumab (1 mg/kg) and anti-SorLA antibody (10 mg/kg). Tumors were monitored and measured every 3-4 days and tumor volume was calculated according to the formula [tumor volume = (L x W x W) x 0.5], where “L” is the length and “W” is the width of xenograft. Mice were sacrificed at day 35 post-engraftment, and tumors were dissected, fixed in 10% formalin, and processed for paraffin sections with standard protocols. Sections were stained for the proliferation marker Ki-67.

### Immunohistochemistry analysis of tumors

Formalin-fixed, paraffin-embedded tissue samples were cut to 4 μm sections, deparaffinized and rehydrated with standard procedures. For immunohistochemistry (IHC) of tumors, heat-mediated antigen retrieval was done in citrate buffer (pH 9). Sections were washed with a 0.05 M Tris-HCl pH 7.6, 0.05 % Tween20 washing buffer, blocked for endogenous hydrogen peroxide activity, and incubated with Normal Antibody Diluent (NABD; Immunologic, #BD09-125). Sections were then incubated with a Ki-67 antibody (Millipore, #AB9260; diluted 1:1000) for 1 h. After washes, samples were incubated for 30 min with a BrightVision Goat anti-Rabbit HRP (Immunologic #DPVR110HRP) secondary antibody, and washed again. After washes, DAB solution (DAKO, #K3468) was added for 10 seconds followed by washing. After counterstain with Mayer’s HTX, slides were dehydrated, cleared in xylene and mounted in Pertex. Stained samples were imaged with Pannoramic P1000 Slide Scanner (3DHISTECH Ltd), and analyzed with QuantCenter software with NuclearQuant quantification module (3DHISTECH Ltd).

### Patient-derived breast cancer explant cultures (PDECs)

Patient tumor samples were collected from consenting patients with permission, approved by the Hospital District of Helsinki and Uusimaa (Ethical permit: 243/13/03/02/2013). A small sample was cut out from a primary tumor obtained from breast cancer surgery and transferred to the laboratory, typically in less than 30 min. Samples were carefully minced with a blade and incubated overnight with gentle shaking (130 rpm) at +37 °C in Mammocult basal medium (StemCell Technologies) containing 0.2% Collagenase A (Sigma), Mammocult proliferation supplements (StemCell Technologies), 4 μg/ml heparin, 0.48 μg/ml hydrocortisone, 50 μg/mL gentamicin and penicillin/streptomycin (all from Sigma). On the following day the mixture was centrifuged at 1300 rcf for 3 min and the pellet was resuspended in 5 ml PBS. Fragments were recentrifuged, resuspended in Matrigel (BD) and seeded in 8-well chamber slides (Nunc) for 3D culture. Samples were cultured in the Mammocult medium described above.

### Immunofluorescence (IF) staining

3D cultured patient samples were fixed with 2% PFA, permeabilized with 0.25% Triton X-100, and blocked with 10% goat serum in PBS. Primary antibodies were diluted in blocking buffer and 3D cultures were stained overnight. After three washes with IF buffer (0.1% BSA, 0.2% Triton X-100, 0.05% Tween-20 in PBS), samples were incubated with secondary antibodies in blocking buffer and washed again. Nuclei were counterstained with DAPI (Thermo Fisher Scientific) and mounted with Immu-Mount reagent (Thermo Fisher Scientific). Imaging was performed using a Leica TCS CARS SP8 confocal microscope (Biomedicum Imaging Unit, University of Helsinki).

### Statistical analyses

At least three independent biological replicates were performed for each experiment. The sample size (N) and the related statistical methods are described within figure legends. When data deviated from a normal distribution based on the D’Agostino-Pearson normality test, non-parametric statistical tests were used. Significance was concluded when a probability value (*P*-value) was lower than 0.05. *NS:* not significant. In every case, unequal variances between groups of data were assumed and two-tailed *P*-values were reported. No power analyses were conducted to estimate sample size.

## Results

### Targeting HER2 downregulates SorLA levels

SorLA promotes HER2 and HER3 oncogenic signaling and exhibits increased expression in brain-trophic metastatic breast cancer cells (18,19). Whether HER2-induced tumorigenesis alters SorLA levels remains unknown. We used the MMTV/C-neu transgenic mice (20) to compare SorLA levels between healthy mammary glands and breast tumors. We found increased SorLA levels in breast tumors (Figure 1A), indicating that HER2-mediated cell transformation increases SorLA expression. Accordingly, HER2 silencing with two independent siRNAs decreased SorLA levels in human HER2-positive (HER2+) BT-474 breast cancer cells (Figure 1B). As an additional control, we targeted HER2 signaling using the dual EGFR/HER2 tyrosine kinase inhibitor lapatinib and assessed SorLA levels in the lapatinib-sensitive HER2+ BT-474 and SK-BR-3 breast cancer cells. Lapatinib treatment was efficient as demonstrated by inhibition of the phosphorylation of mitogen-activated protein kinase 1 and 2 (ERK1/2) (Figure 1C and Figure S1). In line with the siRNA-mediated depletion of HER2, we found that HER2 inhibition with lapatinib decreases SorLA levels in BT-474 and SK-BR-3 cells (Figure 1C and Figure S1). Altogether, these results demonstrate that HER2 expression and kinase activity positively regulate SorLA in breast cancer.

**Figure 1:**
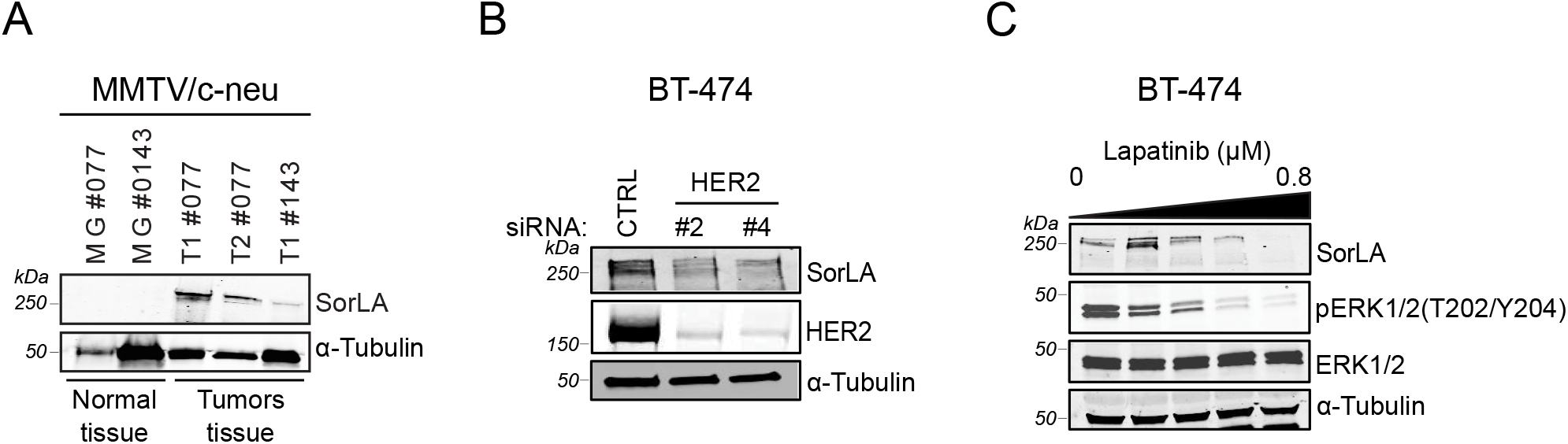
HER2 targeting decreases SorLA levels. A) HER2-driven tumorigenesis increases SorLA levels. Representative immunoblotting of SorLA, with α-tubulin as a loading control, from normal mammary glands (MG#077 and MG#0143) and breast tumors (T1#077, T2#077, and T1#143) from the MMTV/c-neu transgenic mice (20). B) HER2 silencing decreases SorLA levels. Representative immunoblotting of SorLA and HER2, with α-tubulin as a loading control, from HER2-silenced BT-474 cells. C) Lapatinib decreases SorLA levels. BT-474 cells were treated with increasing concentrations of lapatinib (0, 0.1, 0.2, 0.4, and 0.8 μM) for 24 h. Representative immunoblotting of SorLA, pERK1/2(T202/Y204), and total ERK1/2, with α-tubulin as a loading control.

### SorLA silencing blocks S-phase entry and inhibits clonogenic growth

We have previously demonstrated that loss of SorLA inhibits cell viability (18,19), reconfirmed here in BT-474 and MDA-MB-361 cells (Figure S2A and S2B). The effect of SorLA silencing, however, on cell cycle progression remains unknown. Cell cycle analysis using flow cytometry revealed that SorLA silencing triggers an increase and a decrease in the percentage of cells in G1- and S-phase, respectively (Figure 2A). This indicates that SorLA depletion results in a blockade of the cell cycle at G1/S transition, in line with the decreased cyclin D1 expression in SorLA-depleted cells (18). Next, we used our previously established model of breast cancer cells expressing a specific and validated *SORL1* shRNA (18) to investigate the role of SorLA in clonogenic growth. SorLA silencing abolished the ability of single BT-474 and MDA-MB-361 cells to grow into colonies (Figure 2B and Figure S2C). Furthermore, we investigated the effect of SorLA silencing in anchorage-independent growth using the soft-agar colony formation assay. In line with the results of the clonogenic assay, we found that SorLA silencing inhibits colony formation in soft-agar (Figure 2C and 2D). To validate the relevance of targeting SorLA in tumors, we set up CAM assays using HER2-therapy resistant and sensitive MDA-MB-361 and BT-474 cells, respectively (23). Silencing SorLA inhibited tumor growth of both cell lines on the CAM (Figure 2E and 2F; Figure S2D and S2E). Cumulatively, these results demonstrate a role of SorLA in promoting cell cycle progression, cancer cell clonogenicity and tumor growth in HER2-positive breast cancer cells. We aimed next to assess the druggability of SorLA in breast cancer.

**Figure 2:**
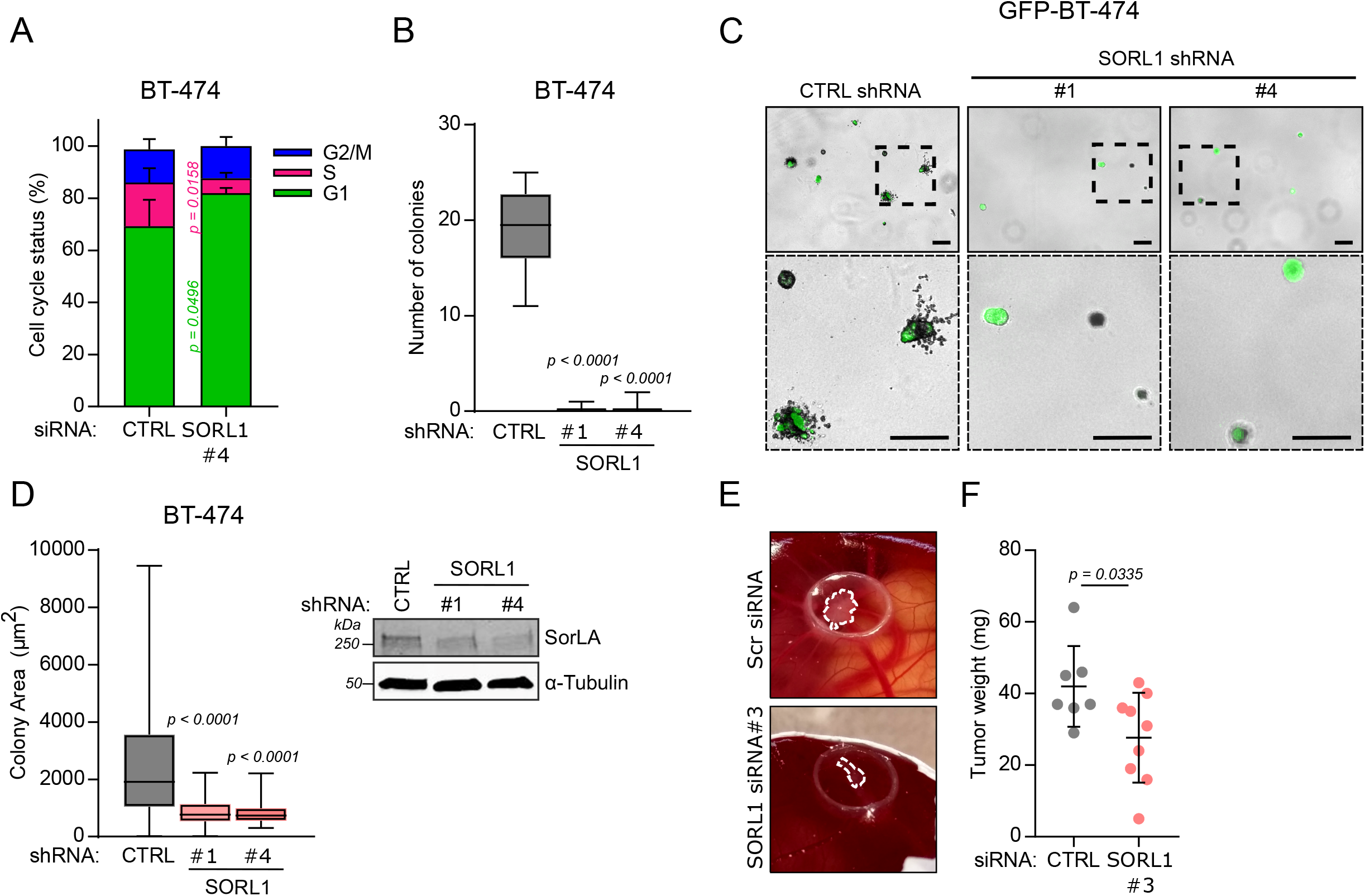
SorLA silencing blocks the cell cycle and inhibits clonogenic growth. A) SorLA silencing inhibits cell cycle S-phase entry. BT-474 cells were silenced for SorLA expression and cells were labeled using the DNA dye propidium iodide (PI) prior to analysis by flow cytometry. Results are represented as mean + SD. B) SorLA silencing inhibits colony formation. Colony formation assay using BT-474 cells stably expressing CTRL shRNA, SORL1 shRNA#1 and shRNA#4. Results are represented as median ± min to max. C) SorLA silencing inhibits colony formation in soft agar. Soft-agar colony formation assay with GFP-expressing CTRL shRNA, SORL1 ShRNA#1 and shRNA#4 BT-474 cells. Scale bars: 100 μm. D) Quantification of colony area from samples described in (C). Results are represented as median ± min to max. N ≥ 20 spheroids / group. A representative western blot validating SorLA silencing is shown. E) SorLA silencing inhibits *in ovo* tumor growth. *In ovo* chorioallantoic membrane (CAM) tumor formation assay with SorLA-silenced MDA-MB-361 cells. F) Tumors described in (E) were weighed and the results are represented as mean ± SD. Statistical analyses: A, Student’s t-test (unpaired, two-tailed, unequal variance); B, Kruskal-Wallis, Dunn’s multiple comparisons test; D, one-way ANOVA, Dunnett’s multiple comparisons test; F, Mann-Whitney U.

### SorLA ab counteracts resistance to HER2-targeted therapy

SorLA is mainly localized at the trans-Golgi network, and undergoes continuous endosomal trafficking between the Golgi and the cell surface (12). Furthermore, we have established that SorLA extracellular domain interacts directly with HER2 and HER3 and that this interaction is necessary for SorLA-dependent proliferation of HER2-positive breast cancer cell lines (24). Thus, we hypothesized that targeting the cell-surface pool of SorLA with a blocking antibody might disrupt its ability to support HER2/HER3 trafficking and thus restrain its oncogenic properties. We chose to investigate a monoclonal SorLA ab that targets the complement-type repeat domains (CRD) in SorLA extracellular part (Figure 3A) as we have established earlier that CRD contributes to the SorLA-HER2 interaction (18). We assessed its ability to inhibit the viability of SorLA silencing sensitive but intrinsically HER2-therapy-resistant MDA-MB-361 cells (23). In line with their resistant background, MDA-MB-361 cells did not respond to HER2-targeting antibody trastuzumab (Figure 3B). Treatment with SorLA ab alone did not exhibit a significant effect on MDA-MB-361 cell viability, indicating that antibody treatment does not have the same effect as depletion of the protein (Figure 3B). However, combining trastuzumab with SorLA ab inhibited the viability of MDA-MB-361 cells (Figure 3B), indicating that SorLA ab alters trastuzumab resistance *in vitro*. Next, we used the CAM assay to validate the growth-inhibitory effect of this combination treatment *in ovo*. Consistent with the cell viability results *in vitro*, we observed that combining SorLA ab and trastuzumab inhibits MDA-MB-361 tumor growth *in ovo* (Figure 3C). In addition, a similar inhibitory effect on tumor growth was observed with a 75% reduced dose of both antibodies (Figure 3C). Altogether, these results demonstrate that, even though SorLA antibody is not sufficient to inhibit cell viability alone, antibody-based targeting of SorLA significantly alters resistance to HER2-targeted therapy *in vitro* and *in ovo*.

**Figure 3:**
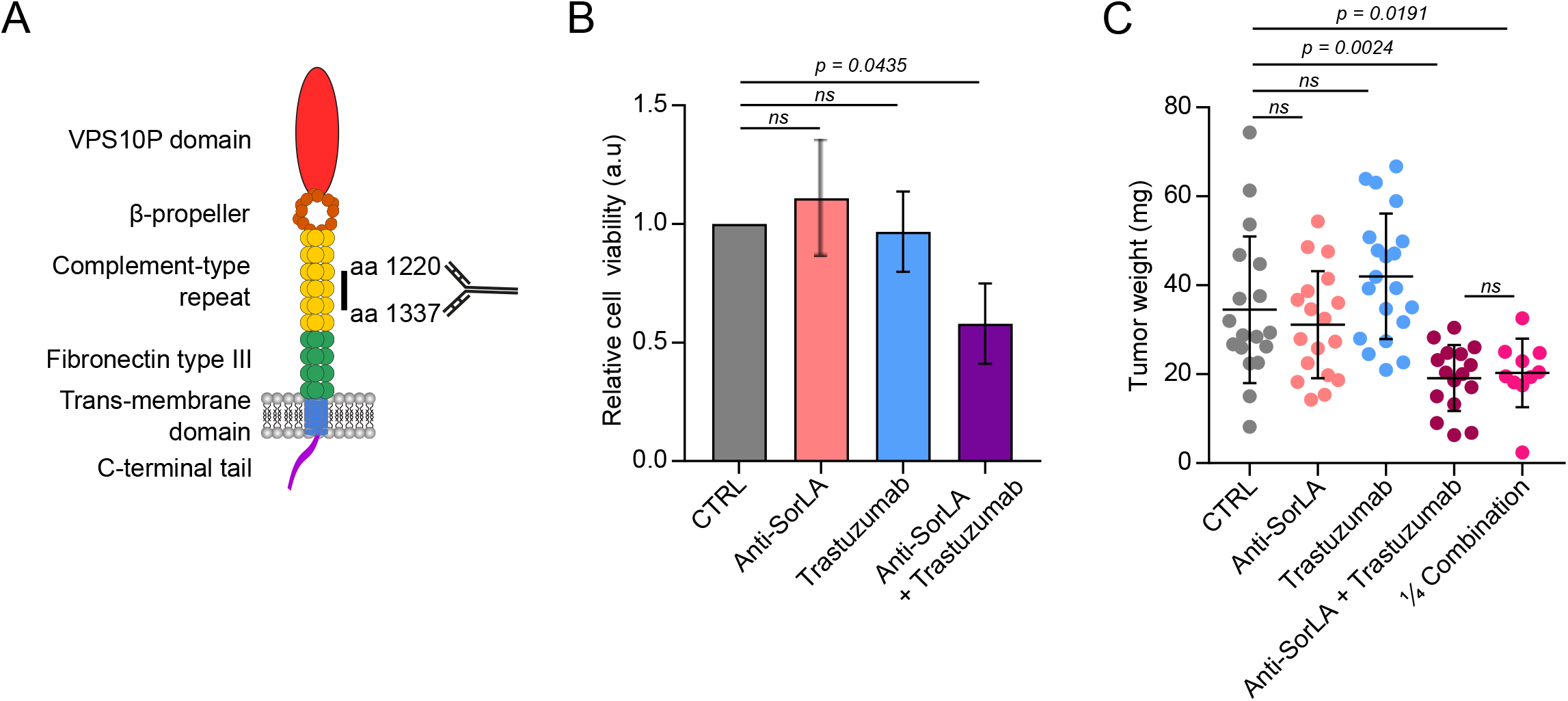
anti-SorLA antibody alters trastuzumab resistance *in vitro* and *in ovo*. (A) Representative scheme of SorLA domains. The epitope in the complement-type repeat domains targeted by anti-SorLA antibody is shown. (B) Anti-SorLA antibody alters the resistance of MDA-MB-361 cells to trastuzumab *in vitro*. MDA-MB-361 cells were treated for 3 days with either IgG control (20 μg.mL^−1^), anti-SorLA antibody (20 μg.mL^−1^), trastuzumab (10 μg.mL^−1^), or a combination of anti-SorLA and trastuzumab. Cell viability was assessed using the WTS-8-based method. Results are represented as mean ± SD. (C) Anti-SorLA antibody alters the resistance of MDA-MB-361 tumors to trastuzumab *in ovo*. MDA-MB-361 cells were engrafted *in ovo* (CAM assay) and treated at days 0 and 3 with 9 μg/10^6^ cells with either IgG control, anti-SorLA antibody, trastuzumab, or a combination of anti-SorLA and trastuzumab and tumors were imaged and weighed at day 5 post-engraftment. ¼ combination corresponds to a 75% reduced concentration of anti-SorLA + trastuzumab. Results are represented as mean ± SD. Statistical analyses: one-way ANOVA, Dunnett’s multiple comparisons test.

### SorLA ab reduces resistance to HER2-targeted therapy in a mouse xenograft model

To investigate the translational relevance of SorLA ab and trastuzumab combination treatment we grafted MDA-MB-361 cells in Nude mice, which subsequently received either IgG control, trastuzumab, or SorLA ab and trastuzumab combination (Figure 4A). Trastuzumab alone slightly inhibited the tumor growth of MDA-MB-361 cells (Figure S3). This effect might have been due to the ability of trastuzumab to trigger a natural killer cell response (25,26). While tumor growth curves revealed no statistical difference between trastuzumab and the combination therapy (Figure S3), further analysis of histopathological features revealed decreased cellularity and lower expression of the proliferative marker Ki-67 in the tumors of mice treated with the combination therapy compared with trastuzumab-treated and control mice (Figure 4B-D). Altogether, these results indicate anti-tumor activity for the SorLA ab and trastuzumab combination treatment *in vivo*.

**Figure 4:**
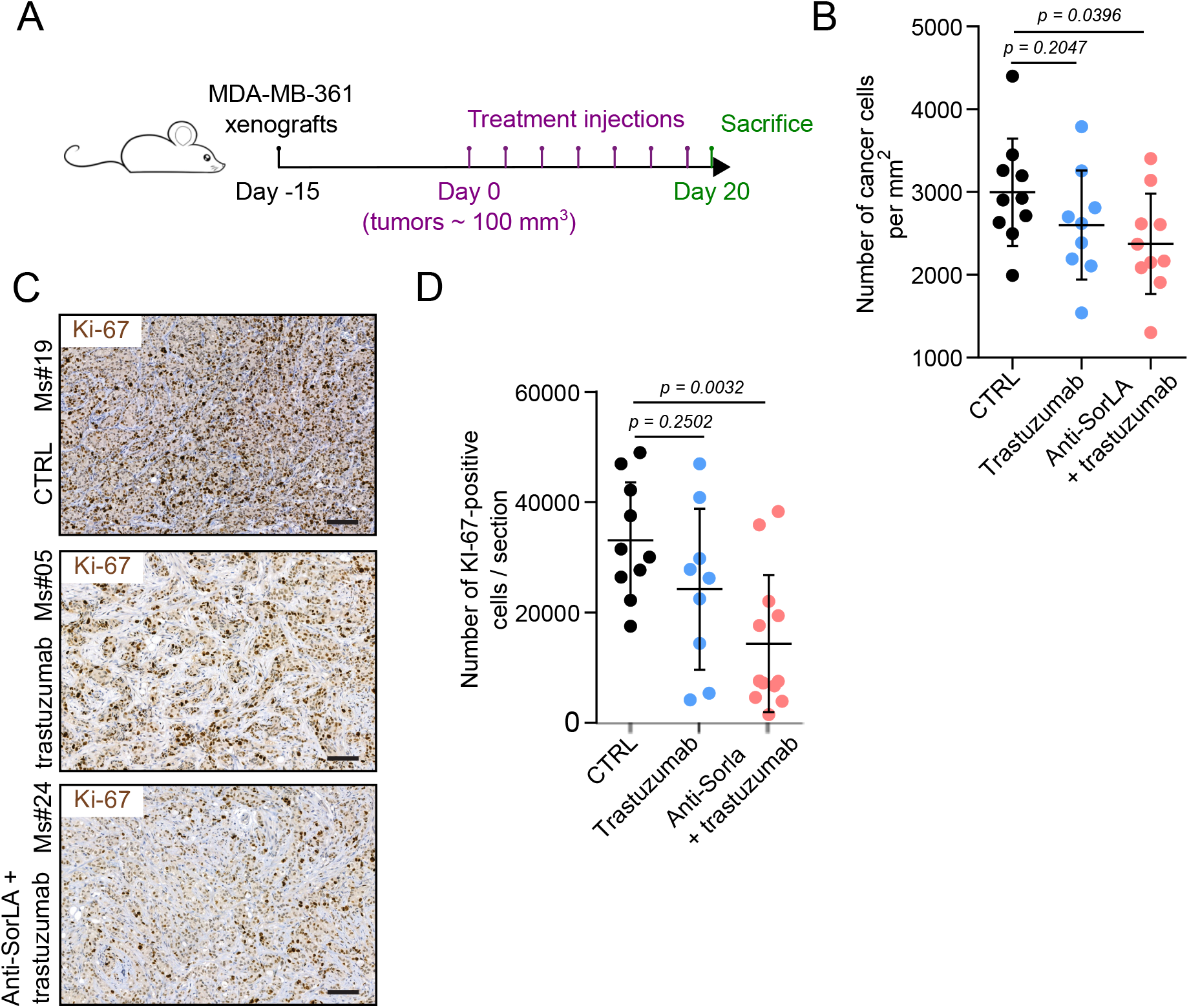
anti-SorLA and trastuzumab combination treatment exhibits antitumor effects *in vivo*. (A) A scheme describing the timeline of the *in vivo* experiment. (B) Anti-SorLA antibody and trastuzumab combination therapy decreases tumor cellularity. Results are represented as mean ± SD. (C) Anti-SorLA antibody and trastuzumab combination therapy decreases the number of Ki-67-positive cancer cells. Tumor sections from IgG control-, trastuzumab-, and anti-SorLA with trastuzumab-treated mice were stained for the proliferative marker Ki-67. Scale bars: 100 μm. (D) Quantification of Ki-67-positive cancer cells in tumor sections described in (C). Results are represented as mean ± SD. Statistical analyses: Student’s t-test (unpaired, two-tailed, unequal variance).

### SorLA ab alters cell proliferation specifically in HER2-positive breast cancer patient-derived explants

To determine whether SorLA ab shows anti-tumor activity in authentic heterogeneous tumor tissue, we explored the effects of SorLA ab in patient-derived explant cultures (PDECs). PDECs were derived fresh from breast cancer surgeries and grown in three-dimensional matrix (27,28). Primary PDECs were treated with SorLA ab or IgG control, and subsequently assessed for the expression of the proliferation marker Ki-67 (Figure 5A). We found that SorLA ab strongly inhibited Ki-67 expression in 2 out of 3 HER2-positive PDECs (Figure 5B). Importantly, SorLA ab did not alter the proliferation of PDECs from HER2-negative breast cancer (Figure 5B), suggesting specific anti-cancer effects for SorLA ab in HER2-positive breast cancer. These results demonstrate that anti-SorLA monoclonal antibody shows activity in a clinically relevant model of HER2-positive breast cancer.

**Figure 5:**
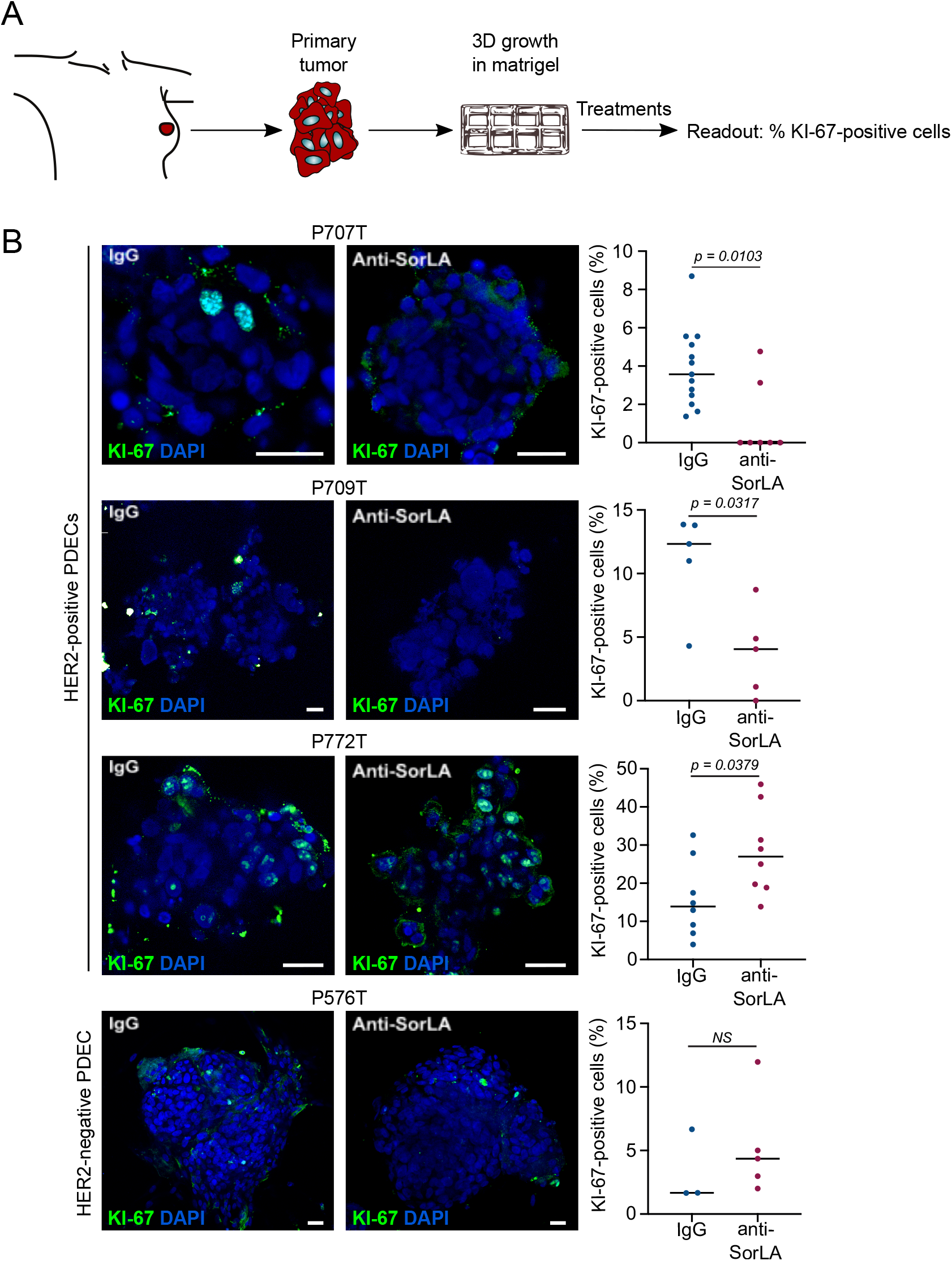
anti-SorLA antibody alters the proliferation of HER2-positive patient tissue-derived explant cultures. (A) Workflow for breast cancer patient tissue-derived explant cultures (PDECs). PDECs were grown in matrigel (see methods for details) and treated with either IgG control or anti-SorLA antibody. Proliferation was assessed using Ki-67 immunofluorescence staining. (B) Anti-SorLA inhibits the expression of the proliferation marker Ki-67 in 2 out of 3 HER2-positive PDECs. PDECs from 3 HER2-positive (P707T, P709T and P772T) and 1 HER2-negative (P576T) patients were treated with either IgG control or anti-SorLA antibody (10 μg.mL^−1^) for 48 h prior to assessment of Ki-67-positive cells by confocal microscopy. Scale bars: 20 μm; each data point represents one organoid; the bars indicate the average percentage of Ki-67-positive cells; statistical analyses: Mann-Whitney U.

## Discussion

Here, we provide for the first time proof-of-concept evidence supporting the druggability of SorLA in cancer. A commercially available SorLA ab altered resistance to HER2 monoclonal antibody trastuzumab *in vitro, in ovo*, and in Nude mouse xenografts *in viv*o. Furthermore, SorLA ab alone exhibited anti-cancer effects specifically in HER2-positive PDECs.

The SorLA extracellular region is a mosaic structure comprising multiple sub-domains shown to mediate ligand binding/discharge. SorLA ab targets the CRD known to be, along with the VPS10P domain, a major site for ligand recognition (29,30) and involved in the direct interaction between HER2/HER3 and SorLA (18,24). We have previously shown that both the extracellular domain of SorLA and its C-terminal tail were necessary for HER2 binding and trafficking, respectively (18). We anticipate that SorLA ab triggers its anti-tumor effects upon binding to the cell surface pool of SorLA prior to the retrograde trafficking pathway since it is reported that low pH, corresponding to acidic milieu of endocytic vesicles, decreases CRD binding to cargo proteins (31).

In our *in vivo* mouse experiment, the combination treatment, but not trastuzumab alone, decreased tumor cellularity and Ki-67 expression, with no difference between tumor growth curves from the combination treatment *vs* trastuzumab. This suggests a delayed action of SorLA ab *in vivo*. A future study could assess the effect of the combination treatment on tumor growth with extended treatment administration. Importantly, we did not observe any signs of toxicity such as weight loss and behavioral changes in mice receiving SorLA ab and trastuzumab combination treatment, albeit the SorLA ab used is reactive against mouse SorLA. This indicates cancer specificity of the treatment most likely owing to the high SorLA overexpression specific for HER2-positive breast cancer cells (18). Furthermore, our CAM assay data shows similar anti-tumor effects between a full and a reduced dose of SorLA ab and trastuzumab combination treatment suggesting that even lower doses of SorLA ab treatment with reduced risk of adverse effects may reach therapeutic efficacy *in vivo*.

Individual SorLA ab treatment was able to inhibit the proliferation in HER2-positive PDECs. These results in a clinically relevant *ex vivo* model, shown to recapitulate the complex tumor heterogeneity (27,32), are promising in terms of translational potential. Interestingly, SorLA ab did not exhibit any significant effects in HER2-negative PDECs. This result is in line with our previous study demonstrating that SorLA promotes oncogenicity specifically in HER2-driven cancers (18).

SorLA promotes bladder cancer growth by sustaining HER2 signaling (18). The efficacy of SorLA ab in inhibiting the progression of SorLA-dependent HER2-driven cancers, other than breast cancer, will need to be assessed in future studies. Furthermore, SorLA overexpression in adipocytes enhances obesity by bolstering insulin receptor signaling through increased receptor recycling (16). Therefore, targeting SorLA in cancer might be translatable to other SorLA-promoted human diseases.

## Supporting information

Al-Akhrass-et-al-Supplementary-Table-and-Figures

## Acknowledgments

We thank J. Siivonen and P. Laasola for technical assistance. H. Hamidi is acknowledged for illustrations and editing the manuscript. The Cell Imaging and Cytometry and Genome Editing core facilities (Turku Bioscience Centre, University of Turku and Åbo Akademi University and Biocenter Finland); the EuroBioimaging node in Turku (Turku Bioscience Centre, University of Turku and Åbo Akademi University) are acknowledged for services, instrumentation, and expertise. This study was supported by an ERC Proof of Concept grant (SaveHER 790274, JI), K. Albin Johansson Foundation (JI), the Finnish Cancer Organizations (JI, JK), the Academy of Finland (JK), Business Finland (JK, PM), EU Horizon 2020 RESCUER (JK), the Finnish Cancer Institute (JI and JK), iCAN Flagship (JK, PM), Jane and Aatos Erkko Foundation (JK) and Sigrid Juselius Foundation (JI, JK). H. Al-Akhrass has been supported by the Finnish Cultural Foundation Central Fund 190150.

## Data availability

The authors declare that the data supporting the findings of this study are available within the article and from the authors on request.

## References

1. Yarden Y, Pines G. The ERBB network: at last, cancer therapy meets systems biology. Nature Reviews Cancer. 2012 Aug;12(8):553–63.

2. Slamon DJ, Leyland-Jones B, Shak S, Fuchs H, Paton V, Bajamonde A, et al. Use of chemotherapy plus a monoclonal antibody against HER2 for metastatic breast cancer that overexpresses HER2. N Engl J Med. 2001 Mar 15;344(11):783–92.

3. Kodack DP, Askoxylakis V, Ferraro GB, Fukumura D, Jain RK. Emerging strategies for treating brain metastases from breast cancer. Cancer Cell. 2015 Feb 9;27(2):163–75.

4. Kodack DP, Chung E, Yamashita H, Incio J, Duyverman AMMJ, Song Y, et al. Combined targeting of HER2 and VEGFR2 for effective treatment of HER2-amplified breast cancer brain metastases. Proc Natl Acad Sci USA. 2012 Nov 6;109(45):E3119–3127.

5. Blackwell KL, Burstein HJ, Storniolo AM, Rugo H, Sledge G, Koehler M, et al. Randomized study of Lapatinib alone or in combination with trastuzumab in women with ErbB2-positive, trastuzumab-refractory metastatic breast cancer. J Clin Oncol. 2010 Mar 1;28(7):1124–30.

6. Geyer CE, Forster J, Lindquist D, Chan S, Romieu CG, Pienkowski T, et al. Lapatinib plus Capecitabine for HER2-Positive Advanced Breast Cancer. New England Journal of Medicine. 2006 Dec 28;355(26):2733–43.

7. Sergina NV, Rausch M, Wang D, Blair J, Hann B, Shokat KM, et al. Escape from HER-family tyrosine kinase inhibitor therapy by the kinase-inactive HER3. Nature. 2007 Jan 25;445(7126):437–41.

8. Garrett JT, Olivares MG, Rinehart C, Granja-Ingram ND, Sánchez V, Chakrabarty A, et al. Transcriptional and posttranslational up-regulation of HER3 (ErbB3) compensates for inhibition of the HER2 tyrosine kinase. Proc Natl Acad Sci USA. 2011 Mar 22;108(12):5021–6.

9. Amin DN, Sergina N, Ahuja D, McMahon M, Blair JA, Wang D, et al. Resiliency and vulnerability in the HER2-HER3 tumorigenic driver. Sci Transl Med. 2010 Jan 27;2(16):16ra7.

10. Mishra R, Patel H, Alanazi S, Yuan L, Garrett JT. HER3 signaling and targeted therapy in cancer. Oncol Rev [Internet]. 2018 May 16 [cited 2020 Mar 10];12(1). Available from: https://www.ncbi.nlm.nih.gov/pmc/articles/PMC6047885/

11. Xie T, Lim SM, Westover KD, Dodge ME, Ercan D, Ficarro SB, et al. Pharmacological Targeting of the Pseudokinase Her3. Nature chemical biology. 2014 Dec;10(12):1006.

12. Willnow TE, Andersen OM. Sorting receptor SORLA – a trafficking path to avoid Alzheimer disease. J Cell Sci. 2013 Jul 1;126(13):2751–60.

13. Fjorback AW, Seaman M, Gustafsen C, Mehmedbasic A, Gokool S, Wu C, et al. Retromer binds the FANSHY sorting motif in SorLA to regulate amyloid precursor protein sorting and processing. J Neurosci. 2012 Jan 25;32(4):1467–80.

14. Jacobsen L, Madsen P, Nielsen MS, Geraerts WPM, Gliemann J, Smit AB, et al. The sorLA cytoplasmic domain interacts with GGA1 and −2 and defines minimum requirements for GGA binding. FEBS Lett. 2002 Jan 30;511(1–3):155–8.

15. Malik AR, Willnow TE. VPS10P Domain Receptors: Sorting Out Brain Health and Disease. Trends in Neurosciences. 2020 Nov 1;43(11):870–85.

16. Schmidt V, Schulz N, Yan X, Schürmann A, Kempa S, Kern M, et al. SORLA facilitates insulin receptor signaling in adipocytes and exacerbates obesity. J Clin Invest. 2016 01;126(7):2706–20.

17. Andersen OM, Reiche J, Schmidt V, Gotthardt M, Spoelgen R, Behlke J, et al. Neuronal sorting protein-related receptor sorLA/LR11 regulates processing of the amyloid precursor protein. Proc Natl Acad Sci USA. 2005 Sep 20;102(38):13461–6.

18. Pietilä M, Sahgal P, Peuhu E, Jäntti NZ, Paatero I, Närvä E, et al. SORLA regulates endosomal trafficking and oncogenic fitness of HER2. Nat Commun. 2019 28;10(1):2340.

19. Al-Akhrass H, Conway JRW, Poulsen ASA, Paatero I, Kaivola J, Padzik A, et al. A feed-forward loop between SorLA and HER3 determines heregulin response and neratinib resistance. bioRxiv. 2020 Jun 10;2020.06.10.143735.

20. Guy CT, Webster MA, Schaller M, Parsons TJ, Cardiff RD, Muller WJ. Expression of the neu protooncogene in the mammary epithelium of transgenic mice induces metastatic disease. Proc Natl Acad Sci U S A. 1992 Nov 15;89(22):10578–82.

21. Laine A, Sihto H, Come C, Rosenfeldt MT, Zwolinska A, Niemelä M, et al. Senescence sensitivity of breast cancer cells is defined by positive feedback loop between CIP2A and E2F1. Cancer Discov. 2013 Feb;3(2):182–97.

22. Schneider CA, Rasband WS, Eliceiri KW. NIH Image to ImageJ: 25 years of image analysis. Nature Methods. 2012 Jul;9(7):671–5.

23. Ding Y, Gong C, Huang D, Chen R, Sui P, Lin KH, et al. Synthetic lethality between HER2 and transaldolase in intrinsically resistant HER2-positive breast cancers. Nat Commun [Internet]. 2018 Oct 15 [cited 2020 Mar 23];9. Available from: https://www.ncbi.nlm.nih.gov/pmc/articles/PMC6189078/

24. Al-Akhrass H, Conway JRW, Poulsen ASA, Paatero I, Kaivola J, Padzik A, et al. A feed-forward loop between SorLA and HER3 determines heregulin response and neratinib resistance. Oncogene. 2021 Jan 8;

25. Arnould L, Gelly M, Penault-Llorca F, Benoit L, Bonnetain F, Migeon C, et al. Trastuzumab-based treatment of HER2-positive breast cancer: an antibody-dependent cellular cytotoxicity mechanism? Br J Cancer. 2006 Jan 30;94(2):259–67.

26. Shi Y, Fan X, Deng H, Brezski RJ, Rycyzyn M, Jordan RE, et al. Trastuzumab Triggers Phagocytic Killing of High HER2 Cancer Cells In Vitro and In Vivo by Interaction with Fcγ Receptors on Macrophages. The Journal of Immunology. 2015 May 1;194(9):4379–86.

27. Turpin R, Munne PM, Suleymanova I, Mustjoki S, Pouwels J, Klefström J. Patient-Derived Explant Cultures (PDECs) as a Model System for Immuno-Oncology Studies. Annals of Oncology. 2019 Dec 1;30:xi45.

28. Haikala HM, Anttila JM, Marques E, Raatikainen T, Ilander M, Hakanen H, et al. Pharmacological reactivation of MYC-dependent apoptosis induces susceptibility to anti-PD-1 immunotherapy. Nat Commun. 2019 Feb 6;10(1):620.

29. Schmidt V, Subkhangulova A, Willnow TE. Sorting receptor SORLA: cellular mechanisms and implications for disease. Cell Mol Life Sci. 2017 Apr 1;74(8):1475–83.

30. Mehmedbasic A, Christensen SK, Nilsson J, Rüetschi U, Gustafsen C, Poulsen ASA, et al. SorLA Complement-type Repeat Domains Protect the Amyloid Precursor Protein against Processing. J Biol Chem. 2015 Feb 6;290(6):3359–76.

31. Caglayan S, Takagi-Niidome S, Liao F, Carlo A-S, Schmidt V, Burgert T, et al. Lysosomal Sorting of Amyloid-β by the SORLA Receptor Is Impaired by a Familial Alzheimer’s Disease Mutation. Science Translational Medicine. 2014 Feb 12;6(223):223ra20–223ra20.

32. Powley IR, Patel M, Miles G, Pringle H, Howells L, Thomas A, et al. Patient-derived explants (PDEs) as a powerful preclinical platform for anti-cancer drug and biomarker discovery. British Journal of Cancer. 2020 Mar;122(6):735–44.

