## Supplementary material for "Sortilin-related receptor is a druggable therapeutic target in breast cancer": Al-Akhrass-et-al-Supplementary-Table-and-Figures

**Supplementary Table: information related to the antibodies used in this study.**

| Antibody | Manufacturer | Catalogue No. |
|---|---|---|
| phospho-ERK p-p44/42 MAPK T202/Y204 | Cell Signaling Technology | #4370S |
| Total ERK p44/p42 MAPK | Cell Signaling Technology | 9102S |
| LR11 (SORL1) | BD Transduction Lab | 612633 |
| HER2/ErbB2 (e2-4001 + 3B5) | Thermo Scientific | MA5-14057 |
| α-tubulin | Hybridoma Bank | 12g10 |
| Ki-67 | Novocastra | ACK02 |

Figure S1

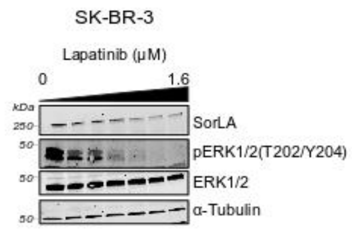


**Figure S1 related to Figure 1**

Lapatinib decreases SorLA levels. SK-BR-3 cells were treated with increasing concentrations of lapatinib (0, 0.1, 0.2, 0.4, 0.8, and 1.6 µM) for 24 h. Representative immunoblotting of SorLA, pERK1/2(T202/Y204), and total ERK1/2, with α-tubulin as a loading control.

Figure S2

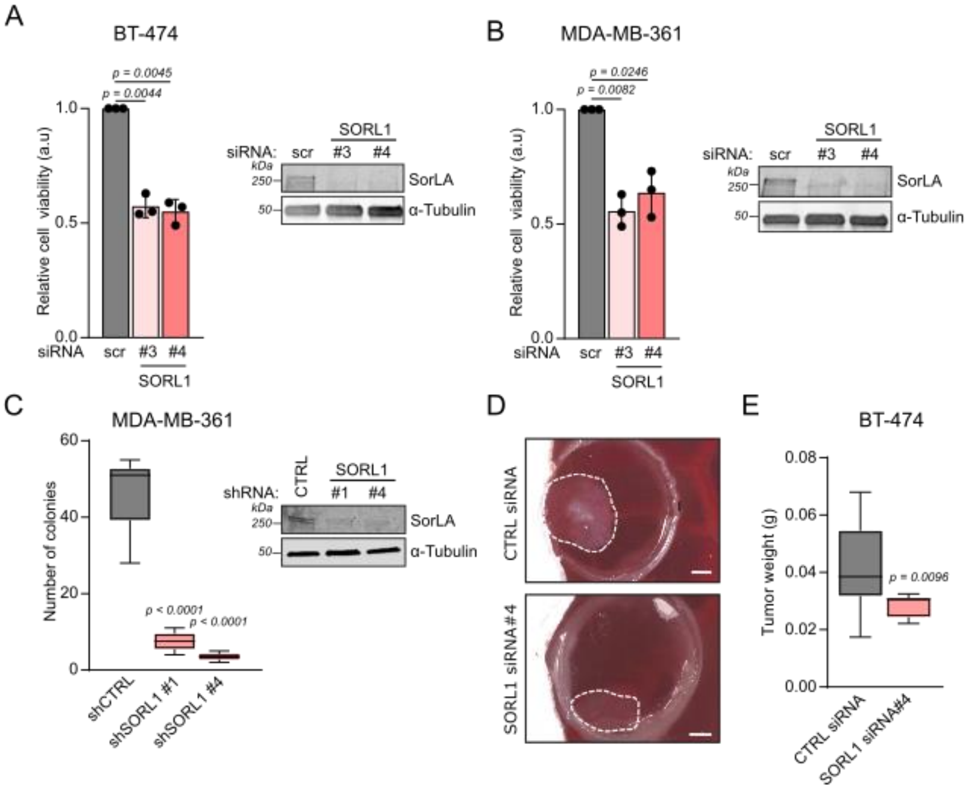

**Figure S2 related to Figure 2**

A) BT-474 cells were transiently silenced for SorLA expression and cell viability was assessed using the WST-8-based method. Each data point represents the average of 3 internal replicates. Results are represented as mean ± SD. A representative western blot validating SorLA silencing is shown.
B) MDA-MB-361 cells were transiently silenced for SorLA expression and cell viability was assessed using the WST8-based method. Each data point represents the average of 3 internal replicates. Results are represented as mean ± SD. A representative western blot validating SorLA silencing is shown.
C) SorLA silencing inhibits colony formation. Colony formation assay using MDA-MB-361 cells stably expressing CTRL shRNA, SORL1 ShRNA#1 and ShRNA#4. Results are represented as mean ± SEM. A representative western blot validating SorLA silencing is shown.
D) SorLA silencing inhibits *in ovo* tumor growth. *In ovo* CAM tumor formation assay with SorLA-silenced BT-474 cells. Scale bars: 1 cm.
E) Tumors described in (D) were weighed and the results are represented as median ± min to max. N ≥ 8 tumors / group.

   Statistical analyses: Student's t-test (unpaired, two-tailed, unequal variance).

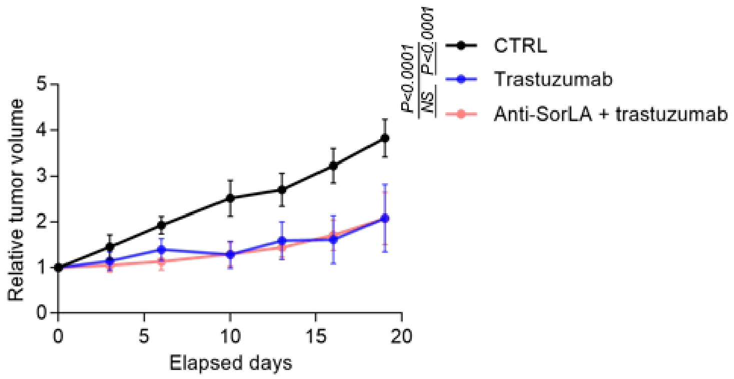


**Figure S3 related to Figure 4**

Tumor growth curves from IgG control-, trastuzumab-, and anti-SorLA with trastuzumab-treated mice. Results are represented as mean relative to day 0 ± SEM. Statistical analyses: exponential growth curve fit comparisons; extra sum-of-squares F test.
